# Image surveys, agricultural landowners and residents hold the key: optimizing local ecological knowledge to reveal carnivore communities

**DOI:** 10.64898/2026.04.25.720805

**Authors:** Elena Fernández-Vizcaíno, Javier Fernández-López

**Author notes:** Corresponding autor.

## Abstract

The choice of appropriate methods to detect species is crucial for biodiversity monitoring. Camera trapping is currently one of the most widely used methods for characterizing mammal communities, although it requires substantial investment in equipment and personnel. In contrast, questionnaires administered to local populations provide a faster and more cost-effective alternative for assessing community composition, but may be influenced by respondent-related biases that compromise data reliability. This study evaluates the concordance between these approaches for characterizing the carnivore community in the Sierra de Segura (Jaén, southern Spain), using Cohen’s kappa coefficient, while also examining the individual and social factors shaping Local Ecological Knowledge (LEK). We deployed 24 camera-trap stations (144 trap nights) across a 25 km^2^ area to record carnivore presence. In parallel, we conducted two types of surveys with local residents (n = 103): (i) free-listing and (ii) image-based species recognition, while recording individual and social characteristics of respondents. Free-listing surveys tended to underreport species, whereas image-based surveys showed higher agreement with camera-trap data, although occasionally overestimating species presence. Higher concordance was associated with social factors indicative of closer and prolonged contact with the environment, such as permanent residence and ownership of agricultural land. Mammal communities differed between methods; however, agreement improved when respondents had higher LEK, while species-specific behavioral traits could also influence perception. Our findings demonstrate that image-based questionnaires can provide results comparable to camera trapping when respondents have strong connections to their natural surroundings. These results highlight the importance of both survey design and respondent selection in improving the accuracy of biodiversity monitoring, offering a transferable framework for integrating LEK into conservation protocols across diverse ecosystems.

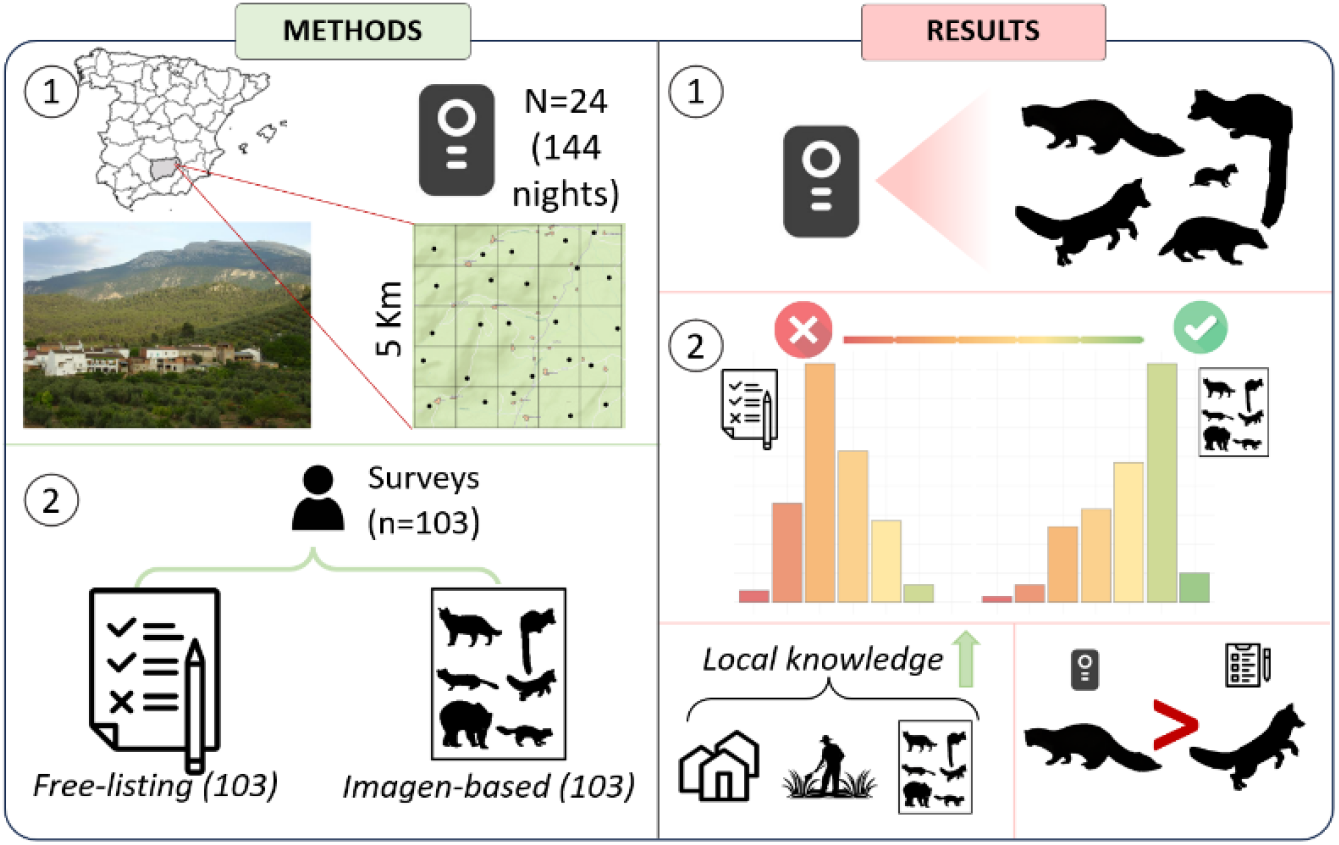

## 1. INTRODUCTION

Camera trapping has become a widely used, non-invasive tool for studying mammal communities due to its versatility across ecological contexts (McCallum, 2013; Steenweg et al., 2017). It provides robust information on detection frequency as well as daily and seasonal activity patterns (Hofmeester et al., 2021, 2019), making it particularly effective for monitoring species that are difficult to observe directly, such as nocturnal or elusive mammals (Gaynor et al., 2018). Despite these advantages, camera trapping is subject to inherent methodological biases (Brittain et al., 2022). Detection probability varies with species-specific traits, including behavior, body mass and home-range size (Hofmeester et al., 2019; Sereno-Cadierno et al., 2026), as well as environmental factors such as vegetation structure and seasonality. For example, detection rates are generally lower in closed than in open habitats (Hofmeester et al., 2019; Sollmann et al., 2013) and may be particularly low for rare, elusive, or focal species (Wong et al., 2019). Moreover, camera trapping can be financially demanding, as achieving reliable results often requires substantial investment in equipment, intensive fieldwork, and extensive data processing, which can limit its applicability under constrained monitoring budgets (Brittain et al., 2022; Gálvez et al., 2016).

As an alternative to camera trapping, survey-based approaches have become increasingly common in wildlife research due to their rapid implementation and cost-effectiveness (White et al., 2005). Surveys have proven particularly useful for documenting mammal species presence (Rodriguez and Delibes, 1992; Burt et al., 2021; Cano and Tellería, 2013; Turvey et al., 2013) and, when appropriately designed, can also provide insights into stakeholder perceptions, human–wildlife interactions, and interdisciplinary conservation strategies (White et al., 2005). Consequently, surveys represent a practical option for biodiversity assessments in logistically challenging areas or under constrained monitoring budgets (Rasalto et al., 2010). However, surveys often present several limitations affecting the reliability of the information they generate. These constraints may arise from the ecological traits of the target species, survey design and respondents’ social and cultural backgrounds (White et al., 2005). In particular, detections based on local ecological knowledge are highly observer-dependent and may vary with seasonality or species behaviour (Boakes et al., 2010; Msoffe et al., 2007; Pearse et al., 2015). For example, elusive or nocturnal species are more likely to be underreported, while limited accessibility in remote areas can further reduce detectability (Brittain et al., 2022). Additionally, reliability of survey-based data also can be strongly influenced by its design; to ensure robust and reproducible results, the target population should be clearly defined, hypotheses explicitly stated, and participant selection procedures carefully documented (White et al., 2005). The use of simple, well-structured question-and-answer formats can reduce misinterpretation by both respondents and researchers (Gomm, 2008). For example, free-listing approaches may underestimate carnivore presence because respondents tend to recall only the most salient or memorable species. In contrast, surveys that provide predefined response options, such as species images, generally yield higher and more reliable detection rates than open-ended questionnaires (White et al., 2005; Gomm, 2008). Finally, social and cultural factors can further influence survey outcomes. Respondents’ familiarity with the study area or target species (Caruso et al., 2017), together with regional variation in socioeconomic conditions and cultural relationships with nature, may introduce context-specific biases in survey-based data (Richardson et al., 2025; White et al., 2005). Despite these limitations, involving local communities in research protocols can strengthen connections to local ecosystems and allow conservation initiatives to benefit from local ecological knowledge (Frigerio et al., 2018; Obura et al., 2021; Pocock et al., 2017). Accordingly, the use of local ecological knowledge within citizen science projects has expanded globally (Hsing et al., 2022; Parsons et al., 2018).

In light of the above, while local ecological knowledge can provide valuable insights, it may also generate unreliable or biased information, highlighting the need for validation against independent field-based data (Can and Togan, 2009). Site-specific evaluations are essential to identify the most effective monitoring approaches under particular ecological and social conditions (Brittain et al., 2022; Zwerts et al., 2021). In human-dominated landscapes, several studies have explicitly compared the detection performance of camera trapping and local ecological knowledge to inform methodological decisions (Brittain et al., 2022; Caruso et al., 2017; Schaller et al., 2012; Zwerts et al., 2021). Ultimately, the optimal approach depends on the target species, available resources, population metrics of interest, and the ecological knowledge of respondents—a factor that is particularly relevant in Spain, where connectedness with nature is among the lowest in Europe (Richardson et al., 2025).

In this context, understanding how ecological, social and methodological factors influence the accuracy of interview based wildlife data is essential for assessing the potential of surveys as a complementary tool for fauna monitoring. The aim of this study is to assess how the kind of survey and social factors influence the concordance between interview-based local ecological knowledge and camera-trap data for monitoring carnivore communities in Mediterranean human dominated landscapes. To do this, we compare information derived from surveys with independent camera trap data and analyze how different survey formats and social factors shape respondents’ knowledge. Specifically, we evaluate the performance of two survey formats, free listing and image recognition, in detecting carnivore species and examine the influence of social variables on the reliability of interview reports in small villages of the Sierra de Segura, Spain. By addressing these questions, this study provides insight into whether and under which conditions interview-based data can serve as a reliable alternative to camera trapping for wildlife monitoring.

## 2. MATERIAL AND METOHDS

### 2. Study areas y camera trapping

The study was conducted in the Natural Park of Sierras de Cazorla, Segura y Las Villas, within a 25 km^2^ area encompassing the municipalities of Orcera and Segura de la Sierra (Jaén, Spain; Fig. 1). The landscape is characterized by Mediterranean pine forests mixed with traditional olive groves, small rural settlements, and agricultural plots such as vegetable gardens and orchards (Herrera, 1998). The study area was subdivided into 25 grid cells of 1 km^2^. One camera trap was deployed per grid cell for six consecutive days, operating continuously (24 h day^−1^). From January to June 2020, between two and six camera traps were active simultaneously and systematically rotated among grid cells, resulting in a total sampling effort of 144 trap-nights. To enhance carnivore detectability and improve community characterization, a bait consisting of a can of sardines was placed approximately 1.5 m in front of each camera (Fernández-López et al., 2014). In total, 24 baited camera trap stations were evenly distributed across the study area. All images were manually reviewed, and species detection–non detection data were extracted to characterize the carnivore community. Independent detection events were defined as consecutive records of the same species separated by at least 10 minutes.

**Figure 1.**
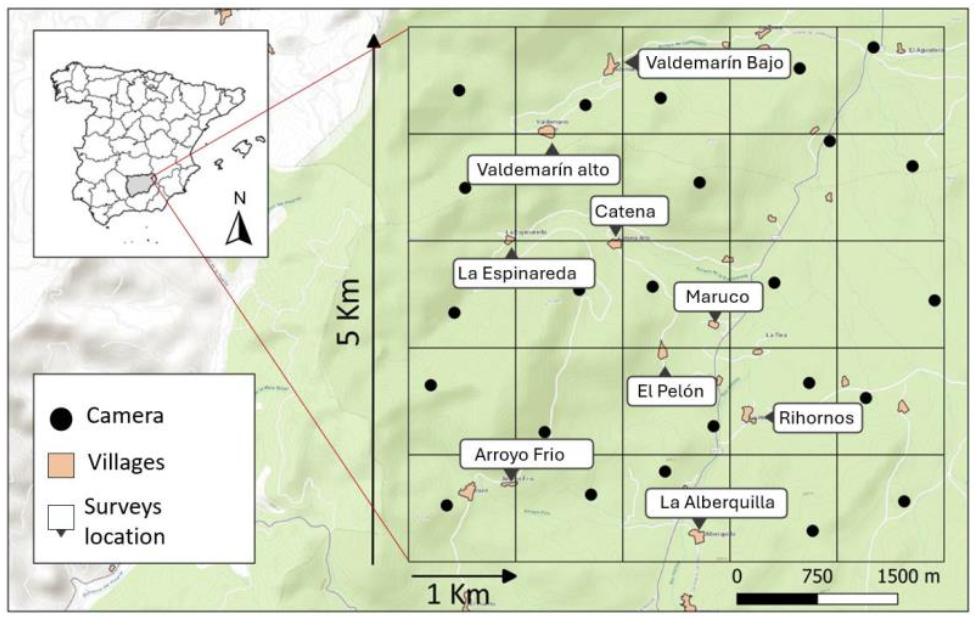
Study area in the municipalities of Orcera and Segura de la Sierra (Jaén, Spain), showing the location of camera trap stations (black dots) and names of villages where interviews were conducted.

### 2. Interview: free-listing survey and image-based survey

Between July and August 2021, two types of face-to-face interviews were conducted with local residents (n = 103) across nine small villages within the 25 km^2^ study area where camera traps were deployed: Valdemarín Bajo, Valdemarín Alto, Catena, La Espinareda, Maruco, El Pelón, Arroyo Frío, Riohornos, and La Alberquilla. Each village has fewer than 100 inhabitants and is located within the municipalities of Orcera and Segura de la Sierra (Jaén, Spain). Participants were contacted in public areas or at their homes. After a brief explanation of the study’s purpose, information was collected to identify potential social factors influencing local ecological knowledge. These factors were assessed through a categorical questionnaire that included age group (18–25, 26–35, 36–45, 46–55, 56–66, >66), gender (male, female, other), permanent residency status (living in the area year-round or for at least three consecutive months per year; yes/no), land ownership (ownership of agricultural land such as olive groves, vegetable gardens, or orchards; yes/no), hunting activity (yes/no), and self-reported knowledge of the natural environment (none, low, medium, high, very high).

Following the collection of sociodemographic information, each participant completed two sections of the interview: a free-listing survey and an image-based species recognition survey. The free-listing survey was always administered first to minimize potential biases arising from visual cues in the image set. Participants were asked to list all wild terrestrial carnivores they knew from the area. In the image-based survey, participants were shown an A3-sized laminated sheet displaying photographs of terrestrial carnivore species known to occur in the Iberian Peninsula, including both native and invasive species, along with their common names (Fig. 2). The species presented were: red fox (*Vulpes vulpes*), stone marten (*Martes foina*), pine marten (*Martes martes*), least weasel (*Mustela nivalis*), European polecat (*Mustela putorius*), Eurasian otter (*Lutra lutra*), common genet (*Genetta genetta*), European badger (*Meles meles*), wildcat (Felis silvestris), Iberian lynx (Lynx pardinus), Iberian wolf (Canis lupus), raccoon (*Procyon lotor*), mink and American mink (*Neovison vison* and *Mustela lutreola*), stoat (*Mustela erminea*), and brown bear (*Ursus arctos*). Images were kindly provided by Dr. José R. Castello from scientific posters and adapted so that species were scaled to reflect their relative body sizes. Presence/absence data for the 16 species were obtained from both survey types at the individual level (free-listing: n = 103; image-based: n = 103). Data were collected anonymously and processed in compliance with Regulation (EU) 2016/679, the General Data Protection Regulation (GDPR). All interviews were conducted in accordance with the ethical guidelines of the Social Research Association (SRA, 2021).

**Figure 2.**
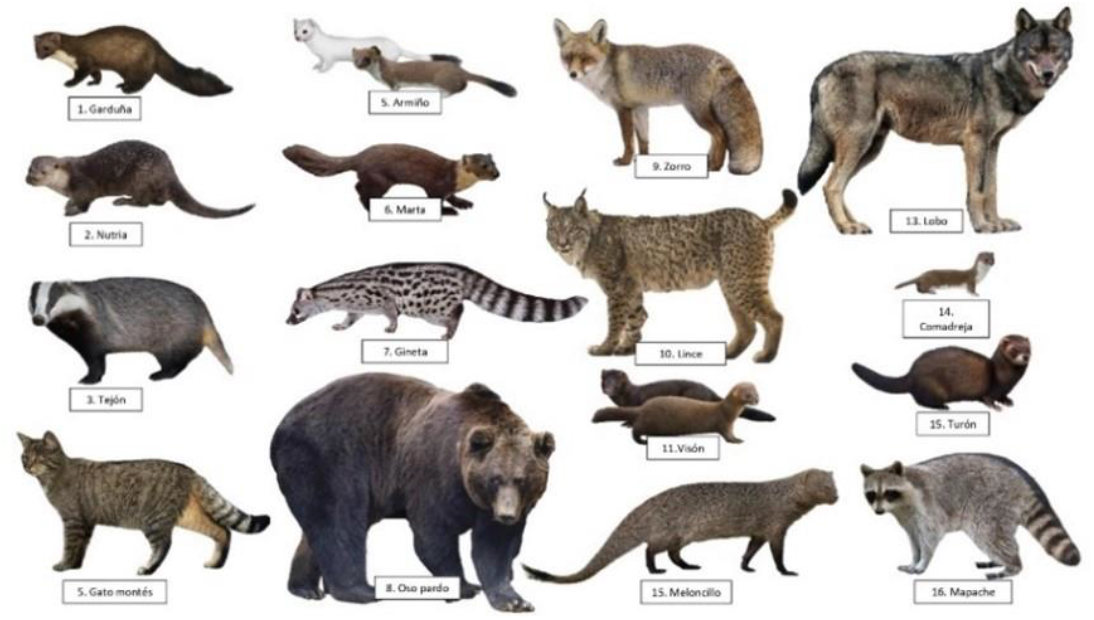
Laminated sheet used in the image-based survey, showing photographs of the 16 wild terrestrial carnivorous mammal species occurring in the Iberian Peninsula, including both native and invasive taxa. Images were provided by Dr. José R. Castello.

### 3. Statistical analysis

First, to evaluate participants’ performance in both the free-listing and image-based surveys, we estimated two descriptive metrics, accuracy and overestimation. Accuracy refers to the proportion of carnivore species correctly identified as present in the study area relative to the total number of species known to occur locally based on camera trap data. Overestimation represents the proportion of species reported by respondents that were not found from camera trapping in the study area. Then, to evaluate the agreement between presence–absence data obtained from camera traps and from the two survey types, we calculated Cohen’s kappa coefficient (Cohen, 1960). Individual kappa values (ranging from −1 to +1) were computed for each respondent, using camera-trap detections as the reference dataset. These coefficients were interpreted as quantitative indicators of local ecological knowledge.

To evaluate the influence of social factors on the level of agreement between interview data and camera trap records (i.e., our knowledge indicator), we used generalized linear mixed models (GLMMs) with a Gaussian distribution. The response variable was the Cohen’s kappa value, and the fixed effects included survey type, gender, age, residence status in the area, land ownership, hunting activity, and environmental knowledge. The village where each interview was conducted was included as a random effect. To further explore how individual and social factors influenced respondents’ knowledge within each survey type, we conducted two additional GLMMs (one for free listing and one for image-based recognition). In these models, Cohen’s kappa values were the response variable, and residency status, hunting activity, environmental knowledge, gender, age, and land ownership were included as predictors, and village was included as a random effect. Initially, all models included all factors and to ensure model robustness, no interaction terms were included. Non-significant terms were subsequently removed through a backward step-selection procedure. Model comparisons were based on the Akaike Information Criterion (AIC), with lower values indicating a better fit. Models differing by at least two AIC units (ΔAIC ≥ 2) with full model were considered substantially better supported.

To explore differences in perceived carnivore community composition among survey groups, we conducted a permutational multivariate analysis of variance (PERMANOVA; Oksanen et al., 2001) based on Jaccard dissimilarities. Because some respondents did not report any species, cases with no species detections were excluded. The model included one fixed factor with three levels: the free-listing survey, the image-based survey, and a subset of image-based respondents who were both landowners and permanent residents. This subset was used to assess whether focusing on the respondent profile identified as most reliable in the GLMMs (i.e., landowners who reside permanently in the area) improves the accuracy of community differentiation. Respondent ID was included in the PERMANOVA to account for repeated measures. All tests were performed with 999 permutations to increase the power and precision of the analysis (Anderson et al., 2008), using residuals under a reduced model (Anderson and Braak, 2003). Pairwise comparisons among survey types were subsequently performed to identify which groups differed significantly in community composition. We also conducted a SIMPER analysis (similarity percentages; Clarke 1993) to determine which species explained the largest proportion of the differences in community composition among study groups. In this study, SIMPER was employed to identify those mammal species responsible for more than 30% of the dissimilarity in each pairwise comparison. To visualize differences in perceived carnivore communities among these groups, we applied a non-metric multidimensional scaling (NMDS) ordination based on Jaccard dissimilarities, using species presence/absence data and stress values were used to assess the goodness of fit of the ordination. All statistical analyses were conducted using R software (R Core Team, 2020) with packages glmmTMB (McGillycuddy et al., 2025) and vegan (Oksanen et al., 2001b). For all tests, the significance level was set at *p* < 0.05. *p*-values between 0.05 and 0.1 were reported as marginally significant effects.

## 3. RESULTS

Camera trap surveys detected five carnivore species: stone marten, red fox, common genet, Eurasian badger, and least weasel, encompassing all species historically known to occur in our study area (Palomo et al., 2007) and achieving a 100% detection efficiency for these focal species. Specifically, stone marten was the most frequently detected species, recorded at 15 out of 24 camera stations with 43 independent detections, followed by red fox at 12 cameras (23 detections), common genet at 11 cameras (17 detections), Eurasian badger at 4 cameras (5 detections), and least weasel with a single detection at one camera station (Table 1). In addition to this five wild species, dogs and cats were also reported in 3 and 1 cameras respectively.

**Table 1.** Occurrence and relative frequency (%) of species, showing their presence in camera traps and across different survey types: free-listing, image-based surveys, and the subset of image surveys from landowners and residents (major ecological knowledge group).

|  | Cameras (N=24) |  |  | Free-listing (N=103) |  |  | Image-based surveys (N=103) |  |  | MRA surveys (N=49) |  |  |
| --- | --- | --- | --- | --- | --- | --- | --- | --- | --- | --- | --- | --- |
| Specie | N | Occurrence (%) | Relative Frequency (%) | N | Occurrence (%) | Relative frequency (%) | N | Occurrence (%) | Relative frequency (%) | N | Occurrence (%) | Relative frequency (%) |
| *Martes foina* | 15 | 62.5 | 34.9 | 31 | 30.1 | 16.8 | 79 | 76.7 | 14.9 | 46 | 93.9 | 15.9 |
| *Vulpes vulpes* | 12 | 50.0 | 27.9 | 80 | 77.7 | 43.2 | 100 | 97.1 | 18.9 | 49 | 100.0 | 16.9 |
| *Genetta genetta* | 11 | 45.8 | 25.6 | 25 | 24.3 | 13.5 | 66 | 64.1 | 12.5 | 44 | 89.8 | 15.2 |
| *Meles meles* | 4 | 16.7 | 9.3 | 12 | 11.7 | 6.4 | 69 | 68.0 | 13.2 | 41 | 82.7 | 14.5 |
| *Mustela nivalis* | 1 | 4.2 | 2.3 | 3 | 2.9 | 1.6 | 60 | 58.3 | 11.3 | 32 | 65.3 | 11.0 |
| *Felis silvestris* | 0 | 0.0 | 0.0 | 16 | 15.5 | 8.6 | 56 | 54.4 | 10.6 | 36 | 73.5 | 12.4 |
| *Lynx pardinus* | 0 | 0.0 | 0.0 | 2 | 1.9 | 1.1 | 15 | 14.6 | 2.8 | 7 | 14.3 | 2.4 |
| *Neovison vison* | 0 | 0.0 | 0.0 | 0 | 0.0 | 0 | 3 | 2.9 | 0.6 | 0 | 0.0 | 0 |
| *Canis lupus* | 0 | 0.0 | 0.0 | 7 | 6.8 | 3.8 | 18 | 17.5 | 3.4 | 5 | 10.2 | 1.7 |
| *Mustela putorius* | 0 | 0.0 | 0.0 | 9 | 8.7 | 4.9 | 26 | 25.2 | 4.9 | 13 | 26.5 | 4.5 |
| *Procyon lotor* | 0 | 0.0 | 0.0 | 0 | 0.0 | 0 | 4 | 3.9 | 0.8 | 1 | 2.0 | 0.3 |
| *Mustela erminea* | 0 | 0.0 | 0.0 | 0 | 0.0 | 0 | 7 | 6.8 | 1.3 | 3 | 6.1 | 1.0 |
| *Lutra lutra* | 0 | 0.0 | 0.0 | 0 | 0.0 | 0 | 18 | 17.5 | 3.4 | 10 | 20.4 | 3.4 |
| *Herpestes ichneumon* | 0 | 0.0 | 0.0 | 0 | 0.0 | 0 | 2 | 1.9 | 0.4 | 1 | 2.0 | 0.3 |
| *Martes martes* | 0 | 0.0 | 0.0 | 0 | 0.0 | 0 | 4 | 3.9 | 0.8 | 0 | 0.0 | 0 |
| *Ursus arctos* | 0 | 0.0 | 0.0 | 0 | 0.0 | 0 | 1 | 1.0 | 0.2 | 1 | 2.0 | 0.3 |

Regarding the surveys, the red fox was the most frequently mentioned species, being detected by 77.75% of respondents through free-listing and 97.1% through the image-based surveys. The stone marten was the second most frequently described species in both surveys, detected by 30.1% of respondents via free-listing and 76.7% via image-based surveys. The common genet ranked third in the free-listing (24.3%) and fourth in the image-based survey (64.4%) while the European badger was mentioned by 11.7% of respondents in the free-listing, placing fourth, and by 69.95% in the image-based survey, ranking third. The least weasel was less frequently mentioned, with 2.95% in the free-listing and 58.3% in the image-based survey. Finally, the wildcat, although not detected by camera traps, was frequently recognized by local knowledge, with 15.5% of respondents mentioning it in the free-listing and 54.4% in the image-based survey. Overall, the patterns of occurrence in detected species are consistent between the two survey methods, with relative frequencies always lower in the free-listing compared to the image-based survey (Table 2; Fig. 5b). This consistency in occurrence is also generally maintained when compared with camera trap detections, except for the red fox, which ranks second in camera trap detections but is the most frequently reported species in both surveys (Table 2; Fig. 5b).

**Table 2.** Mean *Cohen’s Kappa* values (±SD, range) by survey type (*Free-listing* and *Image*), according to *Landownership* and *Residency* status. Bold means marked with an asterisk (*) indicate significantly higher *Kappa* values compared with their respective categories (*p* < 0.05).

|  |  |  | Kappa Cohens values |  |  |  |  |
| --- | --- | --- | --- | --- | --- | --- | --- |
|  |  |  | N | Mean | SD | Min | Max |
| Free listing | Landowner | NO | 37 | 0.22 | 0.17 | -0.12 | 0.67 |
|  |  | YES | 66 | **0.35\*** | 0.23 | -0.12 | 0.85 |
|  | Resident | NO | 48 | 0.24 | 0.19 | -0.12 | 0.67 |
|  |  | YES | 55 | 0.36 | 0.23 | -0.12 | 0.85 |
| Image | Landowner | NO | 37 | 0.42 | 0.25 | -0.12 | 1.00 |
|  |  | YES | 66 | **0.67\*** | 0.21 | -0.05 | 1.00 |
|  | Resident | NO | 48 | 0.46 | 0.29 | -0.12 | 1.00 |
|  |  | YES | 55 | **0.68\*** | 0.16 | -0.05 | 1.00 |

Comparing the detection percentages from surveys with actual presence recorded by camera traps, in the free-listing survey, the mean accuracy rate was 29.3 ± 21.3%, while the mean overestimation rate was 3.0 ± 5.3%. A total of 28.2% of respondents overestimated carnivore presence, reporting on average 10.7% of all carnivore species occurring in the Iberian Peninsula. In the image-based survey, the mean accuracy rate was 73.2 ± 28.6%, and the mean overestimation rate was 13.6 ± 12.5%. In this case, 62.1% of respondents overestimated carnivore presence, with an average overestimation of 18.4% species.

In the general model, Cohen’s kappa values, representing the degree of agreement between species reported in the surveys and those detected by camera traps, were significantly influenced by several factors. Survey type had a strong effect (X^2^ = 81.82, d.f. = 1, p < 0.001; Fig. 3), with image-based surveys showing higher concordance than free-listing surveys. Moreover, permanent residence in the study area (X^2^ = 6.62, d.f. = 1, p = 0.001) and land ownership (χ^2^ = 13.17, d.f. = 1, p < 0.001) were both associated with greater local ecological knowledge, resulting in higher levels of agreement.

**Figure 3.**
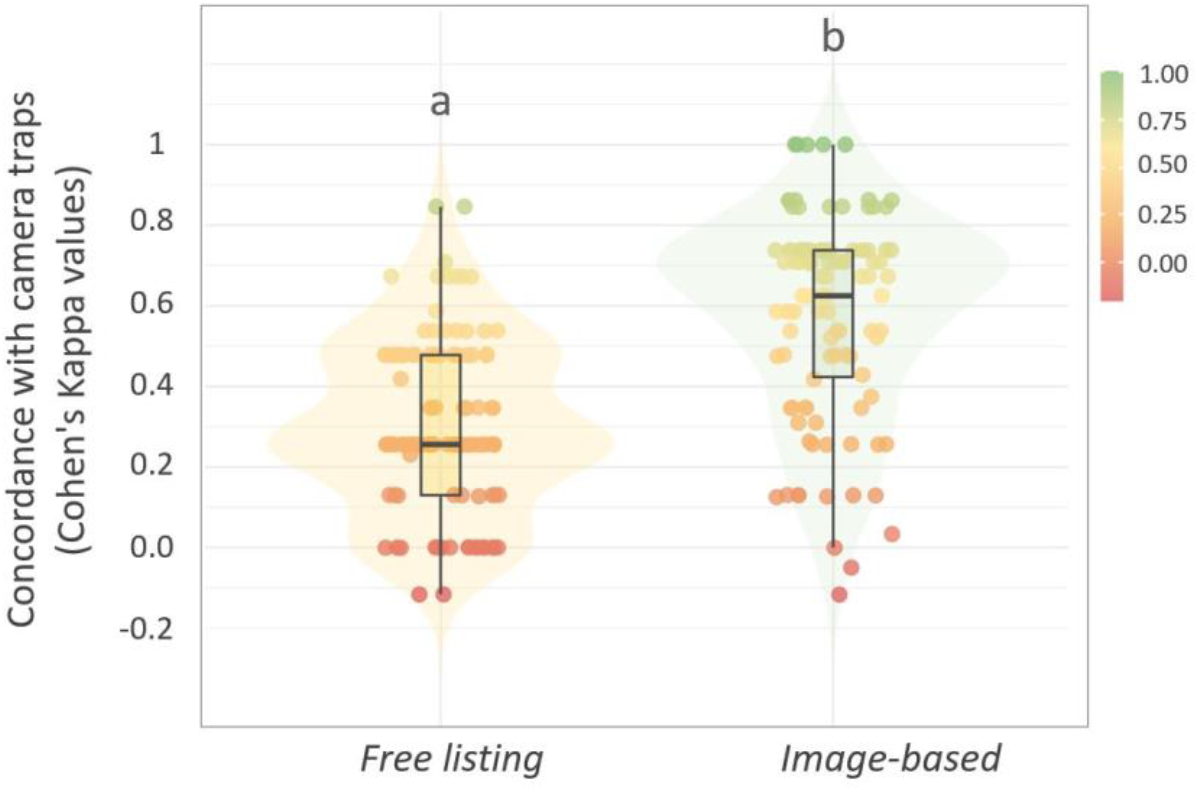
Cohen’s kappa values comparing concordance with camera-trap detections between free-listing interviews (yellow) and image-based interviews (green). Points represent individual respondents, with color intensity reflecting concordance levels. Distributions and central tendencies are summarized using violin plots with overlaid boxplots, and letters indicate significant differences between methods.

**Figure 4.**
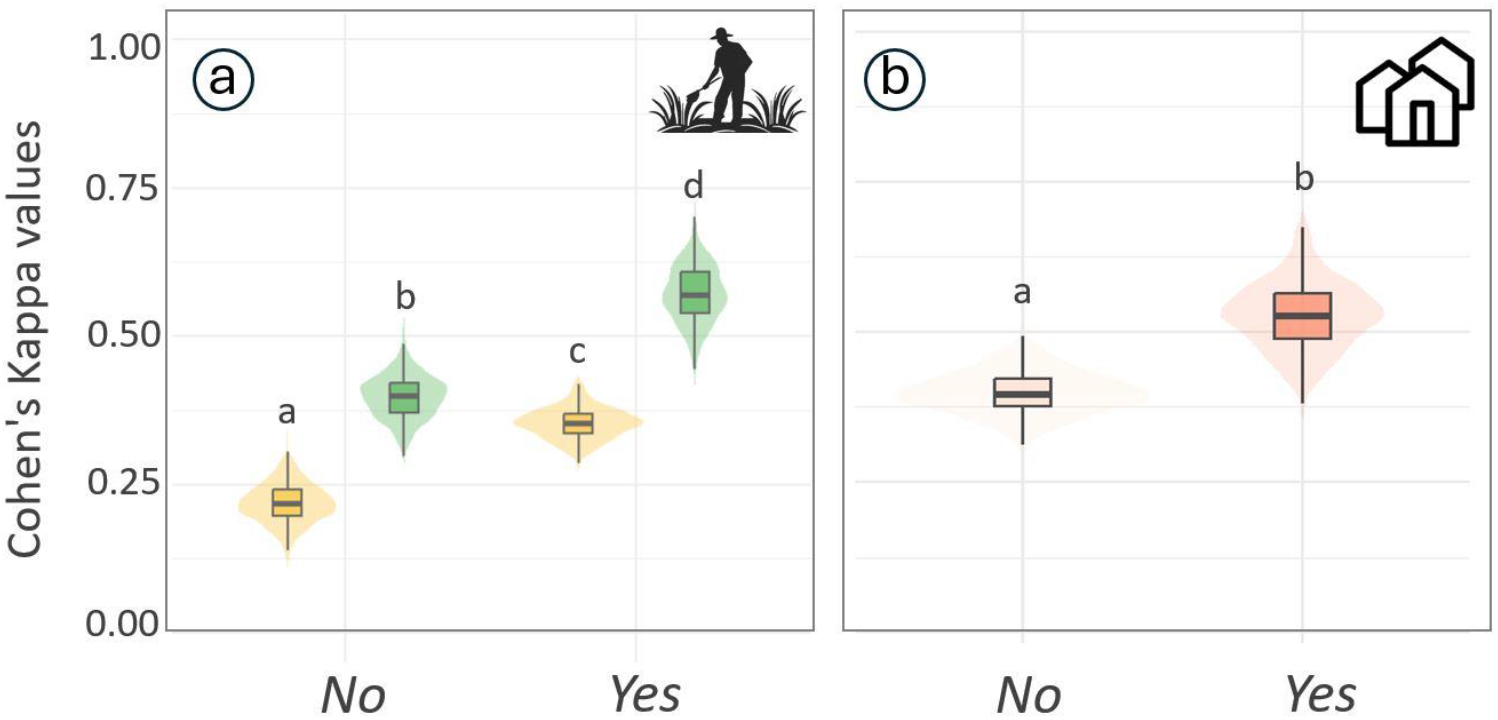
Marginal effects of land ownership and residence status on adjusted Cohen’s kappa values, based on model predictions. Boxplots (±IQR) and violin plots illustrate central tendency and variability around model-predicted means. (a) Effect of agricultural land ownership on adjusted kappa values for free-listing interviews (yellow) and image-based interviews (green). (b) Effect of residence status on adjusted kappa values for image-based interviews. Different letters indicate significant differences between groups (p < 0.05).

In the separate models conducted to further assess which factors influenced each survey type, we found that in the free-listing surveys, agreement levels were influenced only by land ownership, consistent with the trend observed in the general model, with landowners showing higher levels of ecological knowledge (Fig. 3a; X^2^ = 9.95, d.f. = 1, p = 0.002). Conversely, in the image-based surveys, both land ownership (Fig, 3a; X^2^ = 9.86, d.f. = 1, p = 0.002) and permanent residence (Fig. 3b; X^2^ = 9.86, d.f. = 1, p = 0.002) significantly increased the level of agreement, indicating greater knowledge among landowners permanently residing in the area (Table 3). This combination of factors (image-based surveys on land ownerships with permanent residences) was stated as the “*most reliable approach”* (thereafter MRA) for following comparisons.

The PERMANOVA revealed significant differences in perceived carnivore community composition among the different survey types (F = 28.86, R^2^ = 0.20, p = 0.001; Fig 6a). The model explained 19.7% of the total variation in species composition, while the remaining 80.3% was attributed to within-group variation. Pairwise comparisons showed significant differences between all groups. The strongest dissimilarities were found between the free-listing and both the image-based (F = 35.04, R^2^ = 0.16, p = 0.001) and the image survey landowner+resident subsets (F = 44.64, R^2^ = 0.25, p = 0.001). A weaker but still significant difference was detected between the image-based and landowner+resident subsets (F = 3.55, R^2^ = 0.02, p = 0.002). The analysis revealed that no single species explained more than 30% of the dissimilarity by itself; instead, differences were driven by combinations of species. In the comparison between free-listing and image-based surveys, the top five contributing species were badger (8.96%), stone marten (8.71%), genet (7.85%), weasel (7.55%), and wildcat (7.14%), which together accounted for 72.8% of the dissimilarity. For free-listing versus subset MRA surveys, the main contributors were badger (10.49%), genet (8.85%), stone marten (8.61%), wildcat (8.17%), and weasel (7.67%), totaling 78.7% of the dissimilarity. Finally, the difference between image-based and MRA surveys was smaller, with wildcat (4.79%), weasel (4.68%), badger (4.51%), and genet (4.23%) contributing most, for a cumulative contribution of 53.4%. However, significant differences in dispersion among survey types were found (Fig. 5b; F= 12.20, p < 0.001), indicating that PERMANOVA and SIMPER results should be interpreted with caution as group dispersion differs (Fig.6b).

**Figure 5.**
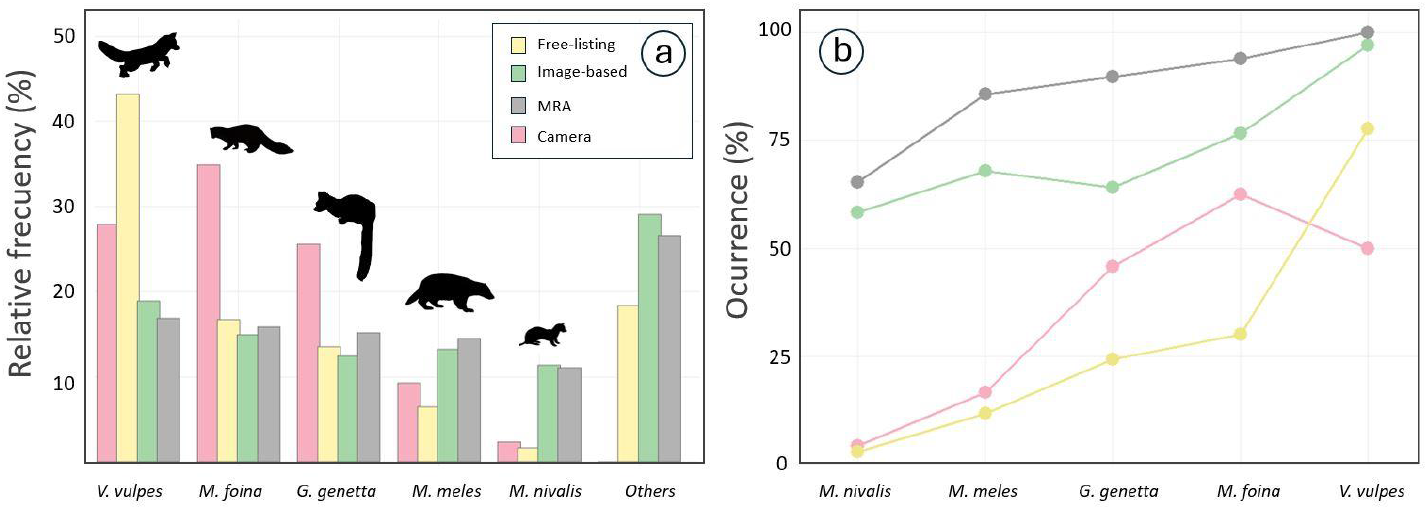
(a) Relative frequency (%) and (b) occurrence of camera or surveys in which each species was detected (%) across survey types: Free listing (yellow), Image based (green), MRA (image+landowner+resident, grey), and Camera (red).

**Figure 6.**
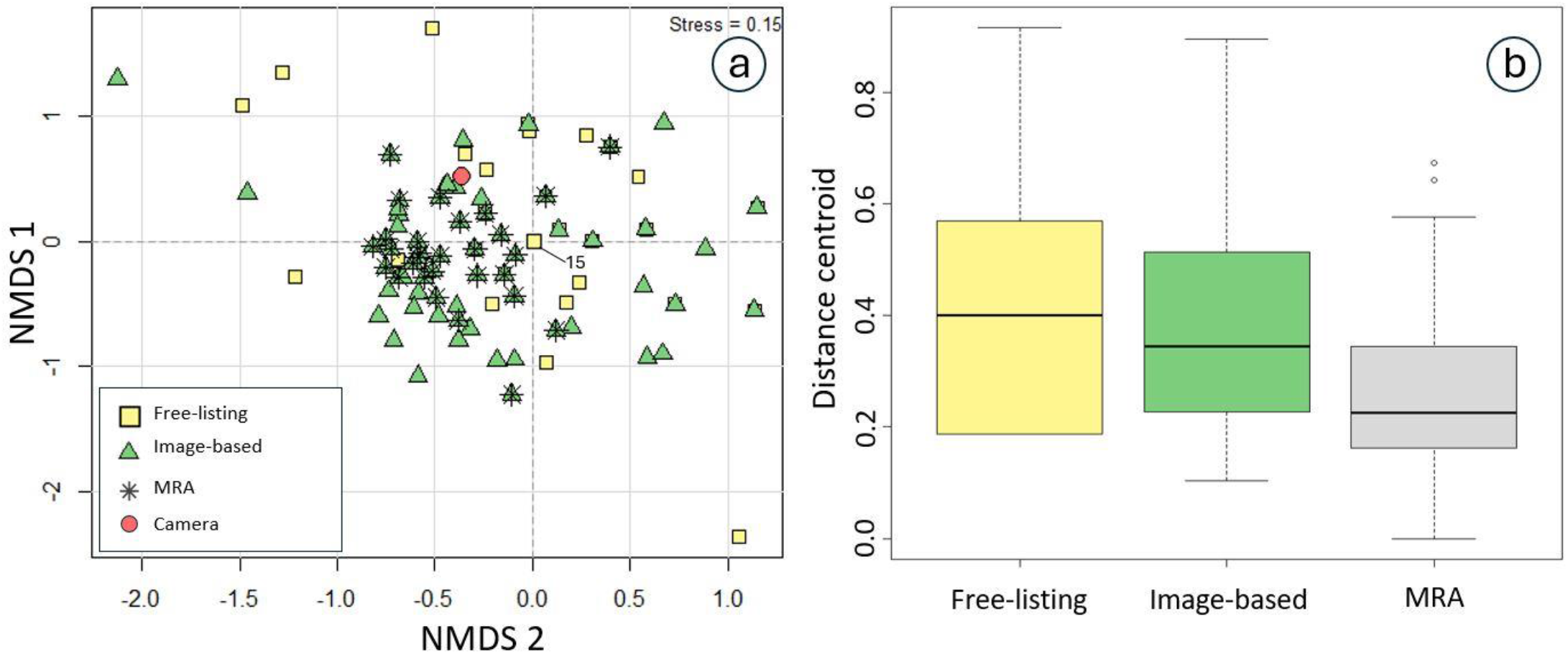
(a) Non-metric multidimensional scaling (NMDS) ordination illustrating differences in perceived carnivore community composition among the three survey groups, free-listing, image-based, and the MRA subset, compared with the community recorded by camera traps.

## 4. DISCUSSION

Our findings support the idea that the type of survey and social factors influence the accuracy of information collected on local carnivore communities through the use of local ecological knowledge. This highlights the importance of evaluating the survey design and target participants prior to implementing a questionnaire. Image-based surveys tend to show higher concordance with camera-trap data, though they often overestimate species presence while free-listing approaches generally underestimate it. Social and contextual factors, such as land ownership and permanent residence in the area, further enhance the concordance between local knowledge and field records. Additionally, respondents’ perceptions of more visible or elusive species can bias report detection and abundance. This indicates that the ecology of the species should be carefully considered. Overall, these results suggest that pre-selecting respondents with specific characters could improve the accuracy and effectiveness of studies using local ecological knowledge for carnivore conservation.

### 4.1. Carnivore community assessment: detectability and perception

Despite our relatively low number of trap-nights (only 144), our camera-trapping results provided a fairly complete picture of the carnivore community previously known from the study area (100% detection; Palomo et al., 2007). Even highly elusive and scarce species such as the least weasel were detected with only six survey days per camera location. This success could be explained by the use of baited camera stations, which may increase detectability of present species as in similar studies (Fernández-López et al., 2014). The use of baited camera traps has been discouraged because baiting may bias animal activity patterns or abundance estimates (Rovero et al., 2013); however, when the objective is limited to detecting species presence, baiting can be a useful tool to increase detection probabilities and optimize sampling effort, as seen in our detection results.

Among the survey questionnaires, the red fox was the most frequently reported species, independently of the survey type (77% in free-listing and 97% in image-based surveys). This result could be explained by its ecological traits: it is more abundant and widespread than other species, shows more diurnal activity, has larger home ranges, and frequently uses human infrastructures such as paths, roads, or human waste as food resources (Díaz-Ruiz et al., 2016; Ferreiro-Arias et al., 2021; Fletcher et al., 2025). However, other factors related to the social perception of this species may also play a role, such as its presence in the media or its involvement in human–wildlife conflicts (e.g. attacks on backyard poultry; van Heel et al., 2017). The common genet and the stone marten were less well known than the red fox, probably due to their more nocturnal activity (Ferreiro-Arias et al., 2021), although their abundance makes them more likely to be detected through indirect observations (e.g. road kills; Grilo et al., 2009). In contrast, the least weasel was almost forgotten by respondents, especially in the free-listing (2.9%), which could be explained by its small size and elusive behavior (King and Powell, 2010; Mos and Hofmeester, 2020). Comparable patterns have also been reported in studies of elusive carnivores: for example, interviews with local inhabitants in Argentina detected the pampas fox (*Pseudalopex gymnocercus*) reliably, while more cryptic species like Geoffroy’s cat (*Leopardus geoffroyi*) and puma (*Puma concolor*) showed low concordance with camera-trap data, highlighting how species traits jointly affect reporting accuracy (Caruso et al., 2017). Additionally, in a study conducted in South Africa comparing camera trap data with online interviews, both methods effectively detected most large mammal species, whereas results were less consistent for smaller species: mammals with low detectability (small-bodied, nocturnal, and elusive) were less likely to be reported by interviewees (Bernard et al., 2024).

A special mention deserves the absence of wildcats in the camera-trap records, despite the species being consistently reported by respondents. This species is currently under research in the Iberian Peninsula, as a recent population decline has been detected (Gil-Sánchez et al., 2020). Our results support previous findings, since although the species was not detected by camera trapping, survey-based information may reflect its presence in nearby zones of our study area where the species has been recently confirmed (SECEM, 2024). The high reporting of this species may also be explained by detections of feral cats in our study, highlighting the importance of considering domestic cats and dogs in camera-trap records. Although their presence was expected given the human-dominated landscape of our study area, it confirms that these species frequently use natural areas in the Iberian Peninsula (Fandos et al., 2012).

### 4.2. Factors shaping local ecological knowledge

Beyond the intrinsic ecological traits of the species, both survey design and respondents’ social characteristics significantly influenced the degree of concordance with camera-trap data, as extensively documented (Berkes et al., 2000; Troudet et al., 2017; White et al., 2005). Survey design showed consistent differences between free-listing and image-based questionnaires: occurrence values were systematically lower in free-listing surveys, indicating generalized underreporting pattern. These results could be explained by recall-related processes, as free-listing requires respondents to retrieve species names from memory without visual or contextual cues, which demands active memory search and greater cognitive effort (Hall et al., 1976; Sousa et al., 2016). In contrast, image-based surveys provide both visual stimuli and written names, reducing memory constraints, facilitating recognition and, therefore, tend to produce higher reporting rates (Hall et al., 1976; Sousa et al., 2016). However, this facilitation effect appears to come at a cost as it also led to higher levels of overestimation, as seen in our results. In this sense, image-based surveys may inflate perceived species richness, particularly for charismatic or well-known species (Troudet et al., 2017). Furthermore, the use of visual stimuli may increase affirmative responses, as participants may feel socially motivated to display knowledge, especially under the perceived evaluation of the interviewer (Nederhof, 1985). Similarly to our findings, another study in insects comparing free-listing surveys with visually supported interviews (photographs and preserved specimens) reported differences between methods, with image stimuli producing more accurate information with field observations (Lima et al., 2016). In that case, free-listing led to underestimation, whereas visual surveys did not result in overestimation because, except for one species, only species present were included. Therefore, in our study, overestimation would likely have been lower had we restricted the list to confirmed carnivore species of the peninsula; however, as this was designed as a methodological test, we intentionally did not control prior knowledge of species presence to assess this potential bias.

Regarding the social factors evaluated, agricultural landownership and permanent residence were the main determinants of local ecological knowledge. Both are associated with greater time spent in direct contact with the environment, which increases opportunities to observe wildlife directly, for example while working the land or through road encounters, and indirectly, such as through predation events affecting livestock. This link between local ecological knowledge, wildlife perception, and the frequency and nature of human–wildlife interactions has been consistently documented in previous studies (Cebrián-Piqueras et al., 2020; Karlsson and Sjöström, 2007). Similarly to our results, previous research in the center of Spain found that local ecological knowledge was strongly related to resident characteristics such as engagement with the natural environment and rural–urban gradient, indicating that individuals who reside in and actively use the landscape develop more detailed ecological knowledge than those less integrated into the local social-ecological context (Cebrián-Piqueras et al., 2020). Díaz-Ruiz et al. (2019), assessing mammal community composition in natural reserve in Chile through free-listing questionnaire surveys also found that the number of species observed was higher in respondents who had lived in the study area for several months than that reported by occasional visitors. Additionally, in our case, the factor *Hunter*, which is also associated with regular contact with nature, did not show significant effects, despite a slight positive trend. In contrast, Descalzo et al. (2023), comparing questionnaires completed by hunters in Spain with data reported by wildlife professionals (researchers and technicians), found high overall concordance but greater Egyptian mongoose presence reported by hunters, suggesting that individuals who spend more time in the field can provide valuable information. However, they are not exempt from biases. This species, for instance, is often perceived by hunters as a threat to small game (Lauret et al., 2026; Martínez-Jauregui et al., 2017), and this negative perception could lead to overreported presence in some cases. In fact, social perceptions of a species, whether positive, neutral, or negative, can influence reported presence and abundance (Troudet et al., 2017; van Heel et al., 2017; Zarazúa-Carbajal et al., 2022), potentially increasing biased responses. Intense public attention to conservation cases, such as the Iberian lynx (Guzmán et al., 2005), as well as to human–wildlife conflicts involving species like the wolf (Karlsson and Sjöström, 2007), can shape social representations of these species and influence how their presence and abundance are reported in surveys, potentially leading to overreporting or misidentification. For example, in the case of the Iberian lynx, several studies reported that interview-based distributions were much broader (Rodríguez and Delibes, 1990) than those confirmed through intensive field surveys, including camera trapping and field signs (Guzmán et al., 2005; Sarmento et al., 2009), suggesting that overreporting may be influenced by the species’ high public visibility and conservation profile, which could have delayed targeted conservation interventions. In our study, although multivariate analyses detected significant differences in community composition between survey types and the subgroup characterized by stronger local ecological knowledge, SIMPER showed that these differences were not driven by any single species with a disproportionately large contribution. This could suggests that variation among surveys was not primarily shaped by species-specific perceptions, but rather reflected broader differences in how communities were described.

Taken together, these findings reinforce that both questionnaire design and respondents’ social characteristics shape biodiversity community assessments based on local ecological knowledge. Survey methods should therefore be aligned with specific conservation objectives. For threatened or conservation-sensitive species, overestimation may generate overly optimistic status assessments, potentially masking population declines and delaying management action, so methods prone to inflation (false positives), such as image-based surveys, should be applied with caution. In contrast, when evaluating, for example, the spread of exotic species, more conservative methods like free-listing may increase the risk of false negatives, delaying early detection and management responses. Therefore, methodological choices should thus balance sensitivity and precaution according to species ecology, social perception, and conservation priorities.

### 4.3. Implications for the use of local ecological knowledge

Our results, together with previous studies, highlight the importance of respondent selection for obtaining reliable biodiversity data (Camino et al., 2020). The PERMANOVA analyses showed that both survey design and social factors significantly shaped the reported community composition, with certain respondent groups producing more consistent species assemblages (Fig. 6). This suggests that controlling for these sources of variation can increase agreement among interviewees and improve the robustness of LEK-based assessments. Testing this methodology in areas with well-documented biodiversity is particularly valuable. While local ecological Knowledge is especially useful in diverse, hard-to-access regions (e.g., tropical forests; Van Vliet et al., 2023), applying it in a well-known system, such as the carnivore community of the Iberian Peninsula, provides a robust reference framework and allows a rigorous evaluation of its performance by comparing survey data with established ecological knowledge, as is the case in our study area.

Our results showed that, based on the frequency of presence reports by respondents, it is possible to establish a threshold above which a species can be assumed to occur in the area. Although this absolute threshold differed between survey types (see table 2; 11%, 58%, and 65% for free-listing, image-based, and MRA surveys, respectively, to detect 80–100% of the species present) the general pattern distinguishing the most and least mentioned species remained consistent across methods, with the exception of the common genet and European badger in image-based surveys. Including multiple species in the survey provides an internal consistency check: if only a single species is assessed, it is harder to evaluate the reliability of responses. By analyzing several species and methods, common reporting patterns (e.g., proportion of respondents and relative ranking) can be identified, providing ancillary information that strengthens conclusions for particular species. Establishing such thresholds has been applied previously; for example, Bernard et al. (2024), in a survey of mammals, only considered species detected by at least 15% of respondents to reduce detection bias and minimize misidentification by excluding outlier responses.

Although questionnaire and interview surveys can sometimes provide reliable estimates of mammal abundance (e.g., Braga-Pereira et al., 2024; Perez-Peña et al., 2012) using species occurrence in surveys as a proxy for abundance is often more challenging (Msoffe et al. 2007; Delibes-Mateos et al. 2014), as noted above, both social and intrinsic species traits influence reporting probability. In our case, when comparing species occurrence in interviews with the proportion of camera traps detecting each species, disagreement was limited to the two most frequently reported species, stone marten and red fox (Fig. 5). Although more detailed studies are needed to directly link these metrics, our results suggest that reporting frequency may broadly reflect relative abundance categories (low, medium, or high) in the surrounding area.

It is important to highlight that our study focuses only on the Order Carnivora, and therefore transferability to other taxonomic groups may be limited, especially because this group is often involved in both cultural and daily human activities. Comparative research should be conducted to evaluate whether the same social factors drive knowledge of less well-known taxa, such as invertebrates or plants. In addition, the size of the study area may limit our findings, since characteristics of the local human population can modulate the applicability of ecological knowledge to inform biodiversity assessments. More structural factors, such as the country of the study area (Spain) and the broader regional context (Western Europe), may also limit the extrapolation of our findings to areas with different levels of nature connectedness 2025 (Richardson et al., 2025). However, this study highlights the importance of previously evaluating the factors affecting the reliability of respondents in order to optimize the use of local ecological knowledge to inform biodiversity status.

## ACKNOWLEDGEMENTS

We are especially grateful to Miguel Delibes-Mateos for his guidance throughout this work and for his valuable feedback during the design of the survey methods. We thank Carmen Vizcaíno Gómez and Manuel Fernández Sánchez for their logistical and field support. We also acknowledge Germán Garrote and Jordi Martinez Guijosa for their assistance with materials. We are additionally grateful to Dr. José R. Castello for providing the images used in the image-based survey. This research did not receive any specific grant from funding agencies in the public, commercial, or not-for-profit sectors.

## REFERENCES

Anderson, M., Braak, C. Ter, 2003. Permutation tests for multi-factorial analysis of variance. J. Stat. Comput. Simul. 73, 85–113. 10.1080/00949650215733

Berkes, F., Colding, J., Folke, C., 2000. Rediscovery of Traditional Ecological Knowledge as Adaptive Management. Ecological Applications 10, 1251. 10.2307/2641280

Bernard, A., Guerbois, C., Venter, J.A., Fritz, H., 2024. Comparing local ecological knowledge with camera trap data to study mammal occurrence in anthropogenic landscapes of the Garden Route Biosphere Reserve. Conserv. Sci. Pract. 6. 10.1111/csp2.13101

Brittain, S., Rowcliffe, M.J., Kentatchime, F., Tudge, S.J., Kamogne-Tagne, C.T., Milner-Gulland, E.J., 2022. Comparing interview methods with camera trap data to inform occupancy models of hunted mammals in forest habitats. Conserv. Sci. Pract. 4. 10.1111/csp2.12637

Burt, C., Fritz, H., Keith, M., Guerbois, C., Venter, J.A., 2021. Assessing different methods for measuring mammal diversity in two southern African arid ecosystems. Mamm. Res. 66, 313–326. 10.1007/s13364-021-00562-x

Camino, M., Thompson, J., Andrade, L., Cortez, S., Matteucci, S.D., Altrichter, M., 2020. Using local ecological knowledge to improve large terrestrial mammal surveys, build local capacity and increase conservation opportunities. Biol. Conserv. 244, 108450. 10.1016/j.biocon.2020.108450

Can, Ö.E., Togan, İ., 2009. Camera trapping of large mammals in Yenice Forest, Turkey: local information versus camera traps. Oryx 43, 427–430. 10.1017/S0030605308000628

Cano, L.S., Tellería, J.L., 2013. Local ecological knowledge as a tool for assessing the status of threatened vertebrates: a case study in Vietnam. Oryx 47, 177–183. 10.1017/S0030605311001669

Caruso, N., Luengos Vidal, E., Guerisoli, M., Lucherini, M., 2017. Carnivore occurrence: do interview-based surveys produce unreliable results? Oryx 51, 240–245. 10.1017/S0030605315001192

Cebrián-Piqueras, M.A., Filyushkina, A., Johnson, D.N., Lo, V.B., López-Rodríguez, M.D., March, H., Oteros-Rozas, E., Peppler-Lisbach, C., Quintas-Soriano, C., Raymond, C.M., Ruiz-Mallén, I., van Riper, C.J., Zinngrebe, Y., Plieninger, T., 2020. Scientific and local ecological knowledge, shaping perceptions towards protected areas and related ecosystem services. Landsc. Ecol. 35, 2549–2567. 10.1007/s10980-020-01107-4

Cohen, J., 1960. A Coefficient of Agreement for Nominal Scales. Educ. Psychol. Meas. 20, 37–46. 10.1177/001316446002000104

Descalzo, E., Ferreras, P., Martínez-Jauregui, M., Soliño, M., Glikman, J.A., Díaz-Ruiz, F., Delibes-Mateos, M., 2023. Assessing the distribution of elusive non-game carnivores: are hunters valuable informants? J. Wildl. Manage. 87. 10.1002/jwmg.22377

Díaz-Ruiz, F., Caro, J., Delibes-Mateos, M., Arroyo, B., Ferreras, P., 2016. Drivers of red fox (*Vulpes vulpes*) daily activity: prey availability, human disturbance or habitat structure? J. Zool. 298, 128–138. 10.1111/jzo.12294

Díaz-Ruiz, F., Caro, J., Ferreras, P., Delibes-Mateos, M., 2019. Assessing mammal community composition in the Huinay Biological Reserve (Chile) through questionnaire surveys: biases associated with respondents. Galemys, Spanish Journal of Mammalogy 31, 1–9. 10.7325/Galemys.2019.A1

Fandos, G., Fernández-López, J., Tellería, J.L., 2012. Incursion of domestic carnivores around urban areas: a test in central Spain. Mammalia 76. 10.1515/mammalia-2011-0050

Fernández-López, J., Fandos, G., Cano, L., García, F.J., Tellería, J.L., 2014. Effect of wildlife refuges on small carnivores in a hunting area in Mediterranean habitat. Hystrix 25, 45–46.

Ferreiro-Arias, I., Isla, J., Jordano, P., Benítez-López, A., 2021. Fine-scale coexistence between Mediterranean mesocarnivores is mediated by spatial, temporal, and trophic resource partitioning. Ecol. Evol. 11, 15520–15533. 10.1002/ece3.8077

Fletcher, J.W.J., Tollington, S., Cox, R., Tolhurst, B.A., Newton, J., McGill, R.A.R., Cropper, P., Berry, N., Illa, K., Scott, D.M., 2025. Utilisation of Anthropogenic Food by Red Foxes (<scp> *Vulpes vulpes* </scp>) in Britain as Determined by Stable Isotope Analysis. Ecol. Evol. 15. 10.1002/ece3.70844

Frigerio, D., Pipek, P., Kimmig, S., Winter, S., Melzheimer, J., Diblíková, L., Wachter, B., Richter, A., 2018. Citizen science and wildlife biology: Synergies and challenges. Ethology 124, 365–377. 10.1111/eth.12746

Gaynor, K.M., Hojnowski, C.E., Carter, N.H., Brashares, J.S., 2018. The influence of human disturbance on wildlife nocturnality. Science (1979). 360, 1232–1235. 10.1126/science.aar7121

Gil-Sánchez, J.M., Barea-Azcón, J.M., Jaramillo, J., Herrera-Sánchez, F.J., Jiménez, J., Virgós, E., 2020. Fragmentation and low density as major conservation challenges for the southernmost populations of the European wildcat. PLoS One 15, e0227708. 10.1371/journal.pone.0227708

Grilo, C., Bissonette, J.A., Santos-Reis, M., 2009. Spatial–temporal patterns in Mediterranean carnivore road casualties: Consequences for mitigation. Biol. Conserv. 142, 301–313. 10.1016/j.biocon.2008.10.026

Guzmán, J., García, F., Garrote, G., Ayala, R., Iglesias, C., 2005. El lince ibérico (Lynx pardinus) en España y Portugal. Censo-diagnóstico de sus poblaciones. Dirección General para la Biodiversidad. Madrid 184.

Hall, J.W., Grossman, L.R., Elwood, K.D., 1976. Differences in encoding for free recall vs. recognition. Mem. Cognit. 4, 507–513. 10.3758/BF03213211

Herrera, C.M., 1998. Flora y fauna del Parque Natural de Cazo ri a, Segura y Las Villas: Presente y futuro de un patrimonio excepcional. Anuario del Adelantamiento de Cazorla 78–83.

Hofmeester, T.R., Cromsigt, J.P.G.M., Odden, J., Andrén, H., Kindberg, J., Linnell, J.D.C., 2019. Framing pictures: A conceptual framework to identify and correct for biases in detection probability of camera traps enabling multi-species comparison. Ecol. Evol. 9, 2320–2336. 10.1002/ece3.4878

Hofmeester, T.R., Thorsen, N.H., Cromsigt, J.P.G.M., Kindberg, J., Andrén, H., Linnell, J.D.C., Odden, J., 2021. Effects of camera-trap placement and number on detection of members of a mammalian assemblage. Ecosphere 12. 10.1002/ecs2.3662

Hsing, P., Hill, R.A., Smith, G.C., Bradley, S., Green, S.E., Kent, V.T., Mason, S.S., Rees, J., Whittingham, M.J., Cokill, J., Stephens, P.A., 2022. Large-scale mammal monitoring: The potential of a citizen science camera-trapping project in the United Kingdom. Ecological Solutions and Evidence 3. 10.1002/2688-8319.12180

Karlsson, J., Sjöström, M., 2007. Human attitudes towards wolves, a matter of distance. Biol. Conserv. 137, 610–616. 10.1016/j.biocon.2007.03.023

King, C.M., Powell, L.A., 2010. The natural history of weasels and stoats: ecology, behavior, and management, 2nd edn. Oxford University Press, Oxford.

Lauret, V., Descalzo, E., Glikman, J.A., Martínez-Jauregui, M., Soliño, M., Ferreras, P., Díaz-Ruiz, F., Delibes-Mateos, M., 2026. Assessing familiarity and potential conflict between hunters and non-hunters regarding mesocarnivore expansion. Biol. Conserv. 313, 111611. 10.1016/j.biocon.2025.111611

Lima, D.C. de O., Ramos, M.A., da Silva, H.C.H., Alves, A.G.C., 2016. Rapid assessment of insect fauna based on local knowledge: comparing ecological and ethnobiological methods. J. Ethnobiol. Ethnomed. 12, 15. 10.1186/s13002-016-0085-z

Martínez-Jauregui, M., Linares, O., Carranza, J., Soliño, M., 2017. Dealing with conflicts between people and colonizing native predator species. Biol. Conserv. 209, 239–244. 10.1016/j.biocon.2017.02.034

McCallum, J., 2013. Changing use of camera traps in mammalian field research: habitats, taxa and study types. Mamm. Rev. 43, 196–206. 10.1111/j.1365-2907.2012.00216.x

McGillycuddy, M., Warton, D.I., Popovic, G., Bolker, B.M., 2025. Parsimoniously Fitting Large Multivariate Random Effects in glmmTMB. J. Stat. Softw. 112. 10.18637/jss.v112.i01

Mos, J., Hofmeester, T.R., 2020. The Mostela: an adjusted camera trapping device as a promising non-invasive tool to study and monitor small mustelids. Mamm. Res. 65, 843–853. 10.1007/s13364-020-00513-y

Nederhof, A.J., 1985. Methods of coping with social desirability bias: A review. Eur. J. Soc. Psychol. 15, 263–280. 10.1002/ejsp.2420150303

Obura, D.O., Katerere, Y., Mayet, M., Kaelo, D., Msweli, S., Mather, K., Harris, J., Louis, M., Kramer, R., Teferi, T., Samoilys, M., Lewis, L., Bennie, A., Kumah, F., Isaacs, M., Nantongo, P., 2021. Integrate biodiversity targets from local to global levels. Science (1979). 373, 746–748. 10.1126/science.abh2234

Oksanen, J., Simpson, G.L., Blanchet, F.G., Kindt, R., Legendre, P., Minchin, P.R., O’Hara, R.B., Solymos, P., Stevens, M.H.H., Szoecs, E., Wagner, H., Barbour, M., Bedward, M., Bolker, B., Borcard, D., Borman, T., Carvalho, G., Chirico, M., De Caceres, M., Durand, S., Evangelista, H.B.A., FitzJohn, R., Friendly, M., Furneaux, B., Hannigan, G., Hill, M.O., Lahti, L., Martino, C., McGlinn, D., Ouellette, M.-H., Ribeiro Cunha, E., Smith, T., Stier, A., Ter Braak, C.J.F., Weedon, J., 2001a. vegan: Community Ecology Package. CRAN: Contributed Packages. 10.32614/CRAN.package.vegan

Oksanen, J., Simpson, G.L., Blanchet, F.G., Kindt, R., Legendre, P., Minchin, P.R., O’Hara, R.B., Solymos, P., Stevens, M.H.H., Szoecs, E., Wagner, H., Barbour, M., Bedward, M., Bolker, B., Borcard, D., Borman, T., Carvalho, G., Chirico, M., De Caceres, M., Durand, S., Evangelista, H.B.A., FitzJohn, R., Friendly, M., Furneaux, B., Hannigan, G., Hill, M.O., Lahti, L., Martino, C., McGlinn, D., Ouellette, M.-H., Ribeiro Cunha, E., Smith, T., Stier, A., Ter Braak, C.J.F., Weedon, J., 2001b. vegan: Community Ecology Package. CRAN: Contributed Packages. 10.32614/CRAN.package.vegan

Palomo, L.J., Gisbert, J., Blanco, J.C., 2007. Atlas y Libro Rojo de los Mamíferos Terrestres de España. Dirección General para la Biodiversidad-SECEM-SECEMU.

Parsons, A.W., Goforth, C., Costello, R., Kays, R., 2018. The value of citizen science for ecological monitoring of mammals. PeerJ 6, e4536. 10.7717/peerj.4536

Pocock, M.J.O., Tweddle, J.C., Savage, J., Robinson, L.D., Roy, H.E., 2017. The diversity and evolution of ecological and environmental citizen science. PLoS One 12, e0172579. 10.1371/journal.pone.0172579

R Core Team (2020), 2020. R: A language and environment for statistical computing. R: A language and environment for statistical computing. R Foundation for Statistical Computing, Vienna, Austria.

Rasalto, E., Maginnity, V., Brunnschweiler, J.M., 2010. Using local ecological knowledge to identify shark river habitats in Fiji (South Pacific). Environ. Conserv. 37, 90–97. 10.1017/S0376892910000317

Richardson, M., Lengieza, M., White, M.P., Tran, U.S., Voracek, M., Stieger, S., Swami, V., 2025. Macro-level determinants of nature connectedness: An exploratory analysis of 61 countries. Ambio. 10.1007/s13280-025-02275-w

Rodríguez, A., Delibes, M., 1990. El lince ibérico (Lynx pardina) en España. Distribución y problemas de conservación. Colección Técnica. Instituto de Conservación de la Naturaleza, Madrid, Spain.

Rovero, F., Zimmermann, F., Berzi, D., Meek, P., 2013. “Which camera trap type and how many do I need?” A review of camera features and study designs for a range of wildlife research applications. Hystrix 24, 148–156.

Sarmento, P., Cruz, J., Monterroso, P., Tarroso, P., Ferreira, C., Negrões, N., Eira, C., 2009. Status survey of the critically endangered Iberian lynx Lynx pardinus in Portugal. Eur. J. Wildl. Res. 55, 247–253. 10.1007/s10344-008-0240-5

SECEM, 2024. Sociedad Española para la Conservación y Estudio de los Mamíferos. Manifiesto sobre la preocupante situación de conservación del gato montés (felis silvestris) en la península ibérica. Sevilla.

Sereno-Cadierno, J., Hofmeester, T.R., Becker, M., Bernard, A., Moolman, L., Fritz, H., Acevedo, P., 2026. Knee height is often right: evaluating device height effects on camera trapping rate. Remote Sens. Ecol. Conserv. 10.1002/rse2.70053

Sollmann, R., Mohamed, A., Samejima, H., Wilting, A., 2013. Risky business or simple solution – Relative abundance indices from camera-trapping. Biol. Conserv. 159, 405–412. 10.1016/j.biocon.2012.12.025

Sousa, D.C.P. de, Soldati, G.T., Monteiro, J.M., Araújo, T.A. de S., Albuquerque, U.P., 2016. Information Retrieval during Free Listing Is Biased by Memory: Evidence from Medicinal Plants. PLoS One 11, e0165838. 10.1371/journal.pone.0165838

SRA, 2021. Social Research Association Research Ethics Guidance. https://the-sra.org.uk/SRA/SRA/Resources/Good-Practice.aspx?hkey=ccb6430d-24a0-4229-8074-637d54e97a5d [accessed March 2021].

Steenweg, R., Hebblewhite, M., Kays, R., Ahumada, J., Fisher, J.T., Burton, C., Townsend, S.E., Carbone, C., Rowcliffe, J.M., Whittington, J., Brodie, J., Royle, J.A., Switalski, A., Clevenger, A.P., Heim, N., Rich, L.N., 2017. Scaling-up camera traps: monitoring the planet’s biodiversity with networks of remote sensors. Front. Ecol. Environ. 15, 26–34. 10.1002/fee.1448

Troudet, J., Grandcolas, P., Blin, A., Vignes-Lebbe, R., Legendre, F., 2017. Taxonomic bias in biodiversity data and societal preferences. Sci. Rep. 7, 9132. 10.1038/s41598-017-09084-6

Turvey, S.T., Risley, C.L., Moore, J.E., Barrett, L.A., Yujiang, H., Xiujiang, Z., Kaiya, Z., Ding, W., 2013. Can local ecological knowledge be used to assess status and extinction drivers in a threatened freshwater cetacean? Biol. Conserv. 157, 352–360. 10.1016/j.biocon.2012.07.016

van Heel, B.F., Boerboom, A.M., Fliervoet, J.M., Lenders, H.J.R., van den Born, R.J.G., 2017. Analysing stakeholders’ perceptions of wolf, lynx and fox in a Dutch riverine area. Biodivers. Conserv. 26, 1723–1743. 10.1007/s10531-017-1329-5

White, P.C.L., Jennings, N.V., Renwick, A.R., Barker, N.H.L., 2005. REVIEW: Questionnaires in ecology: a review of past use and recommendations for best practice. Journal of Applied Ecology 42, 421–430. 10.1111/j.1365-2664.2005.01032.x

Wong, S.T., Belant, J.L., Sollmann, R., Mohamed, A., Niedballa, J., Mathai, J., Street, G.M., Wilting, A., 2019. Influence of body mass, sociality, and movement behavior on improved detection probabilities when using a second camera trap. Glob. Ecol. Conserv. 20, e00791. 10.1016/j.gecco.2019.e00791

Zarazúa-Carbajal, M., Chávez-Gutiérrez, M., Peña-Mondragón, J.L., Casas, A., 2022. Ecological Knowledge and Management of Fauna Among the Mexicatl of the Sierra Negra, México: An Interpretive Approach. Front. Ecol. Evol. 10. 10.3389/fevo.2022.760805

